# Evolution with a seed bank: the population genetic consequences of microbial dormancy

**DOI:** 10.1101/156356

**Authors:** WR Shoemaker, JT Lennon

## Abstract

Dormancy is a bet-hedging strategy that allows organisms to persist through conditions that are sub-optimal for growth and reproduction by entering a reversible state of reduced metabolic activity. Dormancy allows a population to maintain a reservoir of genetic and phenotypic diversity (i.e., a seed bank) that can contribute to the long-term survival of a population. This strategy can be potentially adaptive and has long been of interest to ecologists and evolutionary biologists. However, comparatively little is known about how dormancy influences the fundamental evolutionary forces of genetic drift, mutation, selection, recombination, and gene flow. Here, we investigate how seed banks affect the processes underpinning evolution by reviewing existing theory, implementing novel simulations, and determining how and when dormancy can influence evolution as a population genetic process. We extend our analysis to examine how seed banks can alter macroevolutionary processes, including rates of speciation and extinction. Through the lens of population genetic theory, we can understand the extent that seed banks influence microbial evolutionary dynamics.

## INTRODUCTION

Nature is rarely predictable. Resource availability, disease pressure, and temperature are just a few of the abiotic and biotic factors that fluctuate over space and time. Variation in these, and other factors have important consequences for the growth, survival, and reproduction of individuals. Many taxa respond to variable environmental conditions by entering a reversible state of reduced metabolic activity, a phenomenon known as dormancy (Lennon & Jones, 2011). Dormancy is an adaptive trait that has independently evolved multiple times across the tree of life (Guppy & Withers, 1999). By entering a dormant state, individuals can endure conditions that are suboptimal for growth and reproduction, thereby increasing a population’s long-term geometric fitness (Cohen, 1966). However, dormancy comes at a cost. Not only do dormant individuals miss out on opportunities to reproduce, they must also invest endogenous resources into resting structures and maintenance energy requirements (Bradshaw et al., 1998; Cáceres & Tessier, 2004; van Bodegom, 2007). Despite these costs, dormant individuals accumulate in many systems resulting in the formation of a seed bank (Locey et al., 2017), which serves as a reservoir of genetic and phenotypic diversity (Fig. 1). Seed banks have important implications for a range of ecological processes and patterns, perhaps the most central being the maintenance of biodiversity. Dormancy preserves diversity by reducing interspecific competition allowing for coexistence, a mechanism known as the storage effect (Chesson & Warner, 1981; Chesson, 1994). In addition, seed bank mediated diversity has consequences for other important ecological phenomena including successional dynamics (Marks, 1974; Bazzaz, 1979; Lennon & Jones, 2011), community stability (Kalamees & Zobel, 2002), and ecosystem processes (Wang et al., 2014).

**Figure 1:**
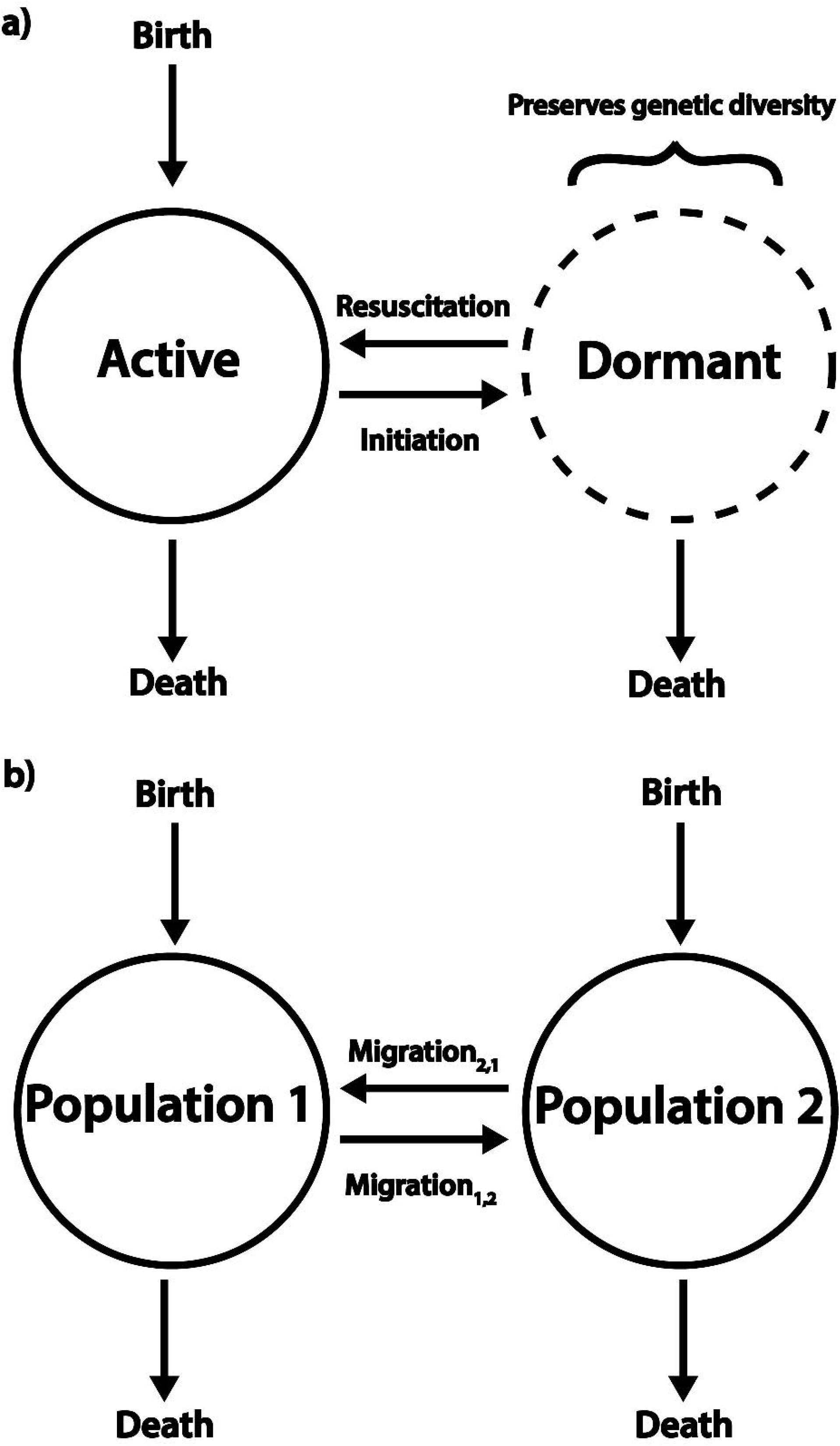
Transitioning in and out of a seed bank is analogous to the migration of individuals between populations. In a) we see a population with a seed bank where individuals switch between active and dormant states. In b) we see two populations where individuals can migrate between each population. An important difference between a) and b) is that individuals must originate in the active pool, because dormant individuals cannot reproduce. While dormant individuals can die in this model, their death rate is much lower than that of active individuals. This conceptual model was first presented in an ecological context (Lennon & Jones, 2011) and later used for the seed bank coalescent (Blath et al., 2015).

Dormancy has important consequences for evolution. For example, rates of phenotypic evolution are reduced in in populations of freshwater zooplankton that are able to persist in a dormant state (Hairston & De Stasio, 1988). Similarly, dormancy is associated with a higher rate of lineage diversification in plants (Willis et al., 2014). However, to fully understand its influence on evolution, it is necessary to understand how dormancy affects the fundamental forces that govern the rate and direction that allele frequencies change over multiple generations. Evolution is a population genetic process governed by both deterministic and stochastic forces, the relative strengths of which dictate how genetic diversity is generated, lost, and maintained. Because dormancy as a life history strategy can provide a fitness benefit, it is subject to the deterministic force of selection (Cohen, 1966; Templeton & Levin, 1979; Brown & Venable, 1986). The ecological implications of dormancy have been extensively studied, particularly the extent that dormancy can outweigh the population genetic effects of alternative life history strategies (Venable & Lawlor, 1980; Venable & Brown, 1988; Olivieri, 2001; Buoro & Carlson, 2014; de Casas et al., 2015). However, far less progress has been made towards understanding how dormancy influences the stochastic forces of evolution (i.e., genetic drift, mutation, and recombination) that operate within a population. To gain a complete understanding of how dormancy influences evolutionary dynamics it is necessary to consider how dormancy affects the stochastic and deterministic forces that underpin evolutionary biology.

While dormancy likely influences eco-evolutionary dynamics for taxa across the tree of life (Hairston et al., 1999; Willis et al., 2014), it could have a particularly large effect on microorganisms, including bacteria, archaea and microeukaryotes, which collectively are, the most abundant and diverse taxa on the planet. Microorganisms have evolved a diverse set of mechanisms that allow individuals to enter and exit a dormant state. These mechanisms include but are not limited to the ability for cells to regulate cellular metabolism, form long-lived endospores, enter a viable but nonculturable state (Oliver, 2005), and produce protective resting stages that are formed during facultative sexual reproduction (Evans and Dennehy, 2005).

Dormancy has attracted attention in the clinical realm because it can help explain how pathogens tolerate high concentrations of antibiotics (i.e., persister cells; Lewis, 2010; Fisher et al., 2017). However, microbial dormancy is also prevalent in complex microbial communities ranging from the human gut to the world’s oceans (Lennon & Jones, 2011). For example, > 90% of microbial biomass is soils is metabolically inactive (Alvarez et al., 1998; Lennon & Jones, 2011; Blagodatskaya & Kuzyakov, 2013).

Microorganisms can quickly transition between active and dormant states (Fig. 2; Walker & Winslow, 1932; Kaprelyants & Kell, 1993; Votyakova et al., 1994; Ishiguro et al., 2015). Although metabolic transition may occur stochastically (Epstein, 2009), dormancy is often regulated by environmental cues, such as changes in temperature (Oliver et al., 1995), pH (Keynan et al., 1964), water (Aadnerud et al., 2015), and resource supply (Dworkin & Shah, 2010). The size and diversity of seed banks can also be affected by the mortality rate of individuals while they are in a dormant state. Some microorganisms succumb to environmental stress within days (Bale et al., 1993), while others can persist in a dormant state for prolonged periods of time. For example, viable microorganisms have been retrieved from ancient materials (e.g., permafrost, amber, halite crystals) that, in some cases, are hundreds of millions of years old (e.g., Johnson et al., 2007). As a consequence, dormant microorganisms can survive for periods of time that far exceed the average generation time of actively reproducing individuals (Cano & Borucki, 1995; Vreeland et al., 2000). Because a considerable fraction of phylogenetically diverse microbial taxa are able to enter a dormant state across disparate environments, it is likely that dormancy has influenced the evolutionary history of microorganisms. Thus, the effects of dormancy have the potential to extend across evolutionary scales, ranging from the population genetic processes that underlie the evolutionary dynamics of populations to the rates that microbial lineages diverge and go extinct.

**Fig. 2.**
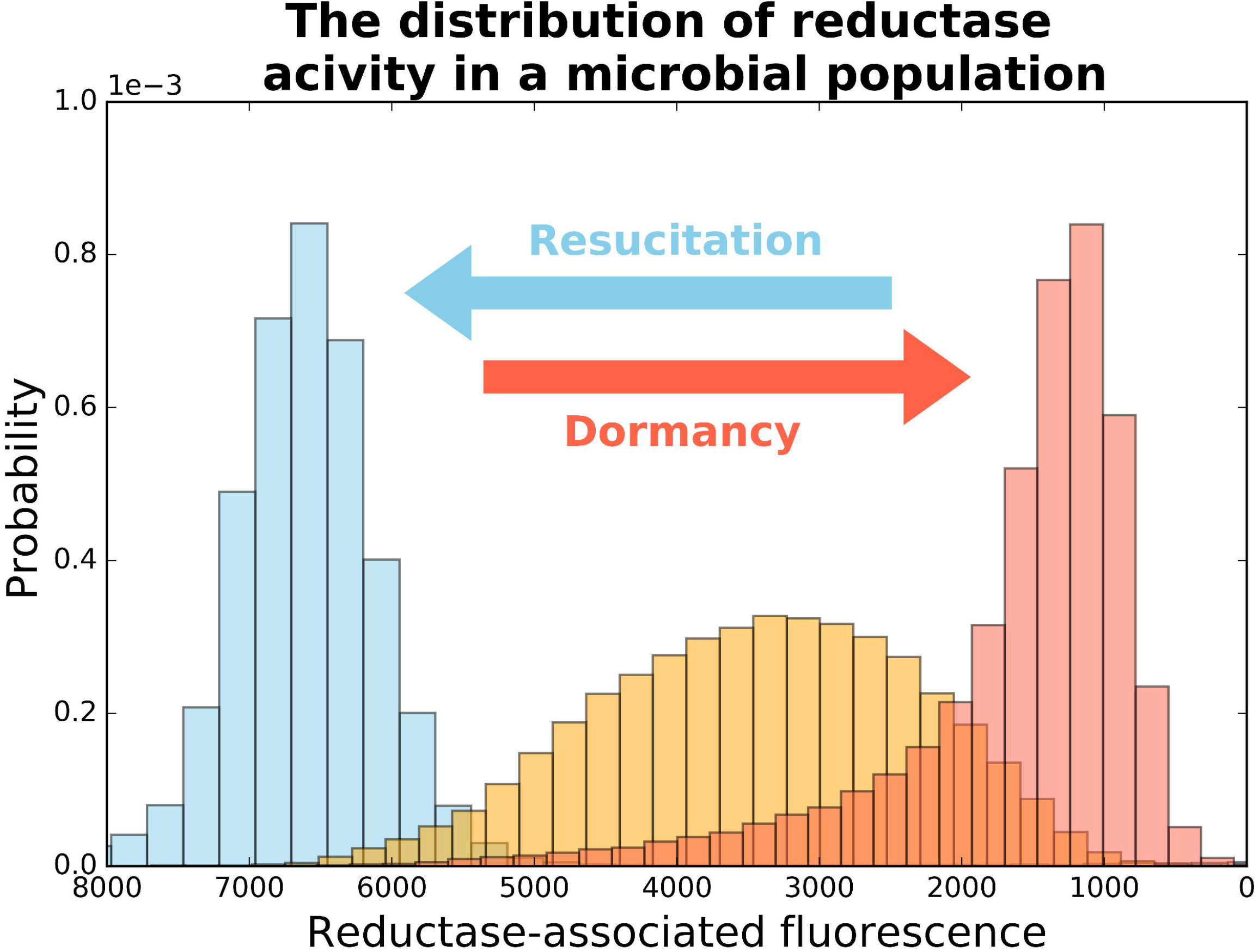
The single cell metabolic activity distribution in a bacterial population at different time-points. The bacterium used is a strain of *Janthinobacterium*, an aerobic, Gram-negative, soildwelling β-proteobacteria. The blue, yellow, and red histograms represent samples taken at 3, 9, and 200 hours after the culture was inoculated, respectively. The distribution of metabolic activity depends on the growth state of the population, where the mode decreases as well as the shape of the distribution as the population enters a dormant state. During the process of resuscitation the mode increases corresponding to an increase in growth rate (see Supplemental Materials).

In this paper, we focus on how dormancy affects microbial evolution. We do this by synthesizing existing population genetic theory and developing novel simulations that provide insight into how seed banks affect the fundamental forces governing the rate and direction of evolution. We then examine how dormancy influences evolutionary dynamics on longer time scales, including macroevolutionary processes such as speciation and extinction. In addition, we examine environments where organisms are often found in a dormant state to determine whether dormancy is an evolutionarily viable bet-hedging strategy (Box 1). This distinction is important because some habitats may not improve on time scales that provide opportunities for reproduction, in which case, dormancy merely increases the amount of time it takes for a population to go extinct. Throughout this paper, we emphasize the importance of using population genetic theory to understand how dormancy as a life history strategy can alter evolutionary dynamics (Box 2). While we focus on dormancy and the evolution of microbial populations, the framework and conclusions should apply to organisms across the tree of life.

## POPULATION GENETIC CONSEQENCES OF DORMANCY

Seed banks preserve genetic and phenotypic diversity by stratifying the population. In a population with a seed bank, there are active and dormant sub-populations, where individuals enter and exit a dormant state in a manner analogous to migration between sub-populations (Lennon & Jones, 2011; Blath et al., 2015) (Fig. 1). Seed banks are sometimes viewed simply as an evolutionary buffer (Koopmann et al., 2016). Dormancy preserves existing genetic diversity by decreasing the rate that genetic diversity is removed from the population (Hairston & De Stasio, 1988; Vitalis et al., 2004; Koopmann et al., 2016), in-turn increasing in the effective size of the population (Nunney, 2002). However, when individuals remain dormant for extremely long periods of time, dormancy can have complex and non-intuitive effects on evolutionary dynamics (Blath et al., 2015). In the following sections, we use population genetic theory to examine how seed banks affect each of the fundamental forces of evolution and expand on that theory through the use of novel simulations.

### Genetic drift

The amount of genetic diversity (*θ*) that can be maintained in an ideal population of finite size is determined by the number of individuals in the population (*N*) and the per-generation mutation rate (*μ*). When the rate that genetic diversity acquired by mutation is equal to the rate that it is lost by drift (i.e., mutation-drift equilibrium), our expectation for the maximum amount of genetic diversity that can be maintained in an ideal population of haploid individuals is *θ* = *2Nμ*. This equation can conveniently be interpreted as the ratio of the rate that genetic diversity is acquired by mutation (*2μ*) and lost by drift (1/*N*) each generation (Kimura, 1969). However, our expectation needs to be modified when a population contains dormant individuals. Because dormant individuals do not reproduce and often have a greatly reduced death rate, genetic diversity turns over at a reduced rate relative to the active portion of the population, reducing the rate of genetic drift, seed banks should increase the maximum amount of genetic diversity that can be maintained in a finite population (Levin, 1990).

Two classes of theoretical population genetic models have been used to explore how genetic diversity scales with dormancy. The first class of models examines what is known as the weak seed-bank effect. The weak seed-bank effect assumes that the maximum number of generations that individuals can spend in an inactive state is smaller than the number of active individuals in the population. Thus, individuals can enter and exit the seed bank before the genetic diversity in the active portion of the population turns over due to the joint effects of mutation and drift (Kaj et al., 2001). The weak seed-bank model predicts that dormant individuals increase genetic diversity, but do not change the pattern of ancestry among individuals in the population (i.e., the shape of the population’s genealogy). In contrast, the strong seed-bank model has no constraint on the maximum number of generations that an individual can remain in a dormant state (González-Casanova et al., 2014; Blath et al., 2015; 2016). Removing this constraint in the mathematical model means that the length of time that an individual spends in the seed bank (where time is measured as generations) can be longer than the number of active individuals in the population. In other words, genetic diversity within the active portion of the population can turn over before a dormant individual is expected to exit the seed bank. Under this scenario genetic diversity within the active portion of the population effectively turns over before an individual in the seed bank has the opportunity to reproduce. The result of the strong seed-bank effect is that dormancy drastically alters the pattern of ancestry among individuals in the total population while increasing the maximum amount of genetic diversity that can be maintained in the population (Fig. 3a). With a strong seed-bank, the expected amount of pair-wise genetic diversity (**E**[π]) in a population with *M* dormant individuals is 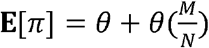. This equation can be interpreted as the amount of genetic diversity expected in an ideal population with *N* actively reproducing individuals (*θ* = *2Nμ*) plus the seed bank’s contribution towards reducing the rate of genetic drift scaled by the relative excess of dormant individuals 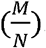. However, it is important to point out that the strong seed-bank effect can require an extremely long period of time to take effect in large populations. For example, assuming a population of *E. coli* of size 10^9^ that is only 10% active (i.e., 10^8^ cells), a strong seed-bank can only occur if on average dormant individuals spend approximately 10^9^ generations in a dormant state. Assuming an average growth rate of ~6.67 generations per day for active individuals (Tenaillon et al., 2016), it would take ~410,000 years for the strong seed-bank to take effect. The length of time required is likely much smaller in real populations, as sub-optimal environmental conditions can reduce the total size of the population and potentially favor dormancy as an adaptive strategy.

**Figure 3:**
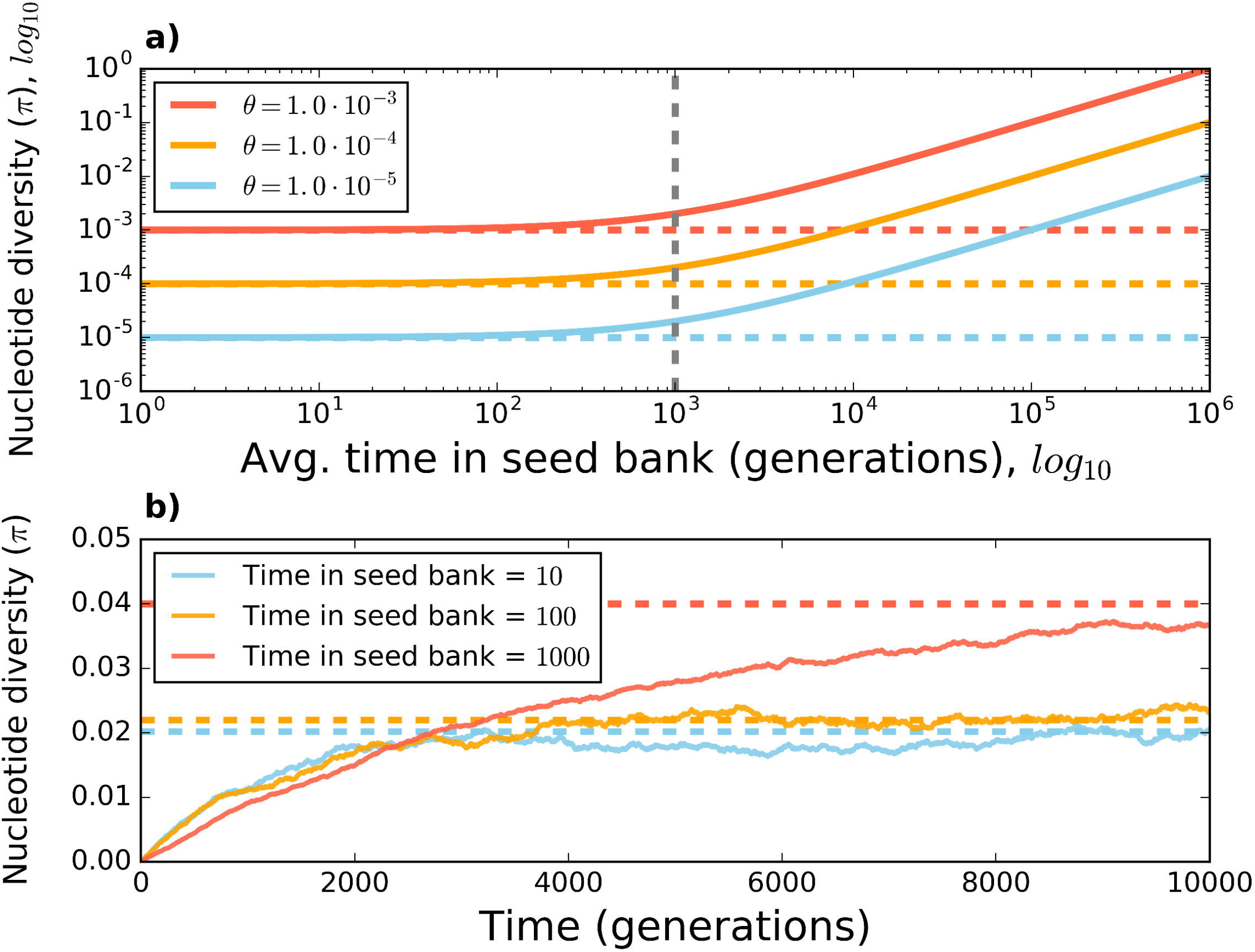
**a)** The expected level of nucleotide diversity (**E**[π]) for a sample of active individuals increases with the number of generations that an individual on average spends in the seed bank for a population at mutation-drift equilibrium with a given population-scaled mutation rate (*θ* - *2Nμ*). Here *N* is the number of active individuals, *M* is the number of dormant individuals, and is the mutation rate for active individuals, where diversity is estimated as 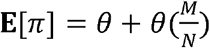 (Blath et al., 2015, eq. 21). The average number of generations that an individual spends in the seed bank can be modeled as a geometric distribution, where the probability of exiting the seed _>_ bank each generation is 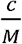 and *c* is the number of individuals that exit the seed bank each generation (Blath et al., 2015; 2016). The vertical grey-dashed line represents the point where the average number of generations that an individual spends in the seed bank is greater than the number of active individuals in the populations (i.e., the strong seed-bank threshold). Vertical lines represent **E**[π] for a population without a seed bank in mutation-drift equilibrium. **b)** The length of time required to reach mutation-drift equilibrium increases with the average number of generations that an individual spends in the seed bank. These results are from a simulated population evolving over time that is subject only to mutation and drift with a constant seed bank size. In both **a)** and **b)** We assume that dormant individuals do not acquire mutations. See Supplementary Information for more detail on implementation of model.

In addition to increasing genetic diversity in a population, our simulations suggest that seed banks increase the average length of time required for a population to reach mutation-drift equilibrium (Fig. 3b). However, while the weak and strong seed-bank models present viable hypotheses about how dormancy can increase genetic diversity, little work has been done to determine if inferred patterns of ancestry in actual populations capable of entering a dormant state resemble predictions from the strong or weak seed-bank effect. Determining whether natural populations capable of entering a dormant state exhibit the strong or weak seed-bank effect will require comparing inferred patterns of ancestry to the predictions of each model.

### Mutation

Mutations are the ultimate source of genetic diversity. In populations composed entirely of reproducing (i.e., active) individuals, the majority of newly acquired genetic variation is due to genome replication errors acquired during cell division (Kunkel, 2004). Given that dormant individuals do not reproduce, it is reasonable to expect that all observed genetic diversity was acquired at some point in the past when the population was active, an assumption that is often made in population genetic models that include dormancy (Vitalis et al., 2004; Blath et al., 2015). However, mutations can still arise in populations that experience sub-optimal conditions. For example, the rate of mutation can effectively be decoupled from the rate of replication in *Mycobacterium tuberculosis* due to oxidative DNA damage that occurs when individuals spend a prolonged period of time in a dormant state (Ford et al., 2011; 2013). This reduced correlation between the rate of replication and the rate of mutation via oxidation-induced DNA damage likely applies to other types of mutagens (e.g., high intracellular concentrations of a chemical or physical mutagen). If acquiring mutations during dormancy is common among microorganisms, then it is necessary to consider the rate of mutation that occurs between replication events as well as the mutation rate during replication to understand how genetic diversity is acquired in populations.

The extent that mutation can be decoupled from replication may depend on the mechanisms regulating transitions into and out of dormancy. For example, DNA is protected during dormancy in endospore-forming taxa like *Bacillus* by the extremely low water content in the endospore coat and small, acid-soluble proteins that bind to the DNA (Setlow, 1992; 2006; Moeller et al., 2009; 2014), as well as the up-regulation of DNA repair during the early phases of resuscitation (Lenhart et al., 2012; Ramírez-Guadianaa et al., 2012). These mechanisms protect endospores from the loss of guanine due to depurination as well as UV radiation-induced pyrimidine dimers (Setlow, 1992), suggesting that environmentally induced mutagenesis occurs at a reduced rate in endospores. However, many taxa enter and exit dormancy without producing specialized resting structures (e.g., endospores, cysts, akinetes). As such, non-sporulating taxa may only rely on mismatch repair mechanisms that are upregulated during resuscitation (Mizrahi & Anderson, 1998; Rittershaus et al., 2013) or DNA repair mechanisms that can be maintained under an energy-limited state (Gong et al., 2005; Dieser et al., 2013; Rittershaus, 2013). For non-endospore-forming taxa that spend extended periods of time in an energy-limited state it is likely that the long-term mutation rate is primarily determined by the fidelity of DNA repair mechanisms that operate during dormancy, rather than that of polymerases used during periods of rapid growth. Maintaining DNA repair may be a better mechanism for long-term cell survival rather than an absolute lack of metabolic activity. For example, evidence of DNA repair has been found in bacteria isolated from half-million-year-old permafrost samples (Johnson et al., 2007).

While the exact cost of maintaining DNA repair mechanisms relative to the total energy budget of energy-limited cells is not known, it is small enough that it has been ignored when estimating the cost of cell maintenance under conditions of homeostasis (Lynch & Marinov, 2015) and extreme energy limitation (Kempes et al., 2017). These assumptions suggest that a certain level of mismatch repair can be maintained over long periods of dormancy without draining the total cellular budget.

Attempting to reproduce under sub-optimal conditions, rather than going dormant, can elevate the rate of mutation in a population. Microbial mutation rates tend to be higher under stressful conditions due to the upregulation of error-prone machinery used for DNA replication and repair (Witkin, 1976; Foster, 2007). Although controversial, it has been argued that this error-prone machinery may be adaptive under times of stress, as an increased mutation rate should increase the number of beneficial mutations acquired over a given length of time (Rosenberg, 2001; Foster, 2007; Galhardo et al., 2007). For example, it has been argued that DNA polymerases used under times of stress can confer a competitive advantage to starved populations of *Escherichia coli*, a phenomenon that has been dubbed Growth Advantage at Stationary Phase (GASP) (Finkel, 2006). However, any temporary fitness advantage due to an upregulated mutation rate will have long-term consequences for the survival of the population. Because almost all mutations are deleterious (Lynch et al., 1999), any increase in the mutation rate will be accompanied by a proportional increase in the average number of deleterious mutations acquired per-individual per-generation, contributing towards the long-term deterioration of the genome (Gerrish et al., 2013; Lynch et al., 2016). The exact amount that fitness is reduced depends on the mutational distribution of fitness effects, which in-turn depends on the environment and the spectrum of mutations generated by the set of molecular machinery used for DNA replication and repair. For example, populations of *E. coli* upregulate DNA polymerase (Pol) IV under times of stress, a highly mutagenic polymerase that is capable of synthesizing DNA across lesions in the genome that would stop alternative polymerases (Ling et al., 2001). Under extreme stress, the ability to use an error-prone polymerase has a clear fitness advantage over death (McHenry, 2011; MacLean et al., 2013). However, alternative error-prone polymerases will almost certainly be accompanied by a proportional increase in the number of deleterious mutations acquired per unit time. The accumulation of deleterious mutations would result in a decrease in fitness once the environment improves, suggesting that populations that continue to grow under a starved state likely constitute an evolutionary dead end. Rather, if there is sufficient temporal variation in the environment then it is possible that the long-term geometric fitness of a population would be maximized by persisting in a dormant state rather than attempting to reproduce and incurring the negative effects of an elevated mutation rate.

### Selection

A large body of theoretical and empirical research has focused on the evolutionary and ecological dynamics that emerge when dormancy is favored by selection (Hedrick, 1995; Nunney, 2002; Malik & Smith, 2008; Ayati & Klapper, 2010). However, far less work has been done to understand how the ability to remain in a dormant state alters the ability for natural selection to act on a population. Because dormancy can act as a buffer against the stochastic forces of mutation and genetic drift, seed banks should reduce the rate that the directional force of natural selection removes genetic diversity from the population. Natural selection is less efficient in models that assume a weak seed-bank effect, where the average amount of time required for selection to drive a beneficial allele to fixation (*T_fix_*) accelerates quadratically with the average number of generations that an individual spends in a dormant state (Koopmann et al., 2016). However, little work has been done to examine selection under a strong seed-bank effect or how selection affects *T_fix_* for active and dormant portions of the population. Intuitively, one might expect that *T_fix_* would increase with the average number of generations that an individual spends in a dormant state, since individuals can spend an even longer period of time in a dormant state. Consequently, beneficial alleles should take longer to go to fixation because individuals must enter and exit the seed bank to spread the beneficial allele throughout the whole population. To test this hypothesis, we simulated the trajectory of a beneficial allele in a population subject to the strong seed-bank effect and examined the dormant and active portions of the population separately (see Supplemental Materials: Selection simulation). As expected, we found that the average amount of time required for a beneficial allele to reach fixation increases with the average number of generations that an individual spends in the seed bank (Fig. 4a, b) (Koopmann et al., 2016).

However, because any fitness advantage only increases the frequency of a beneficial allele if its carrier is reproducing, it is necessary to examine the active sub-population separately from the dormant sub-population. In our model, the beneficial allele initially rose in frequency until it was effectively fixed (a.k.a., quasi-fixation), where it then fluctuated below a frequency of one as individuals continued to exit and enter the seed bank, in a fashion analogous to back-mutation in a single-locus model. By examining the active portion of a population with a seed bank in a simulation of selection on a single locus, we found that the average number of generations required for a beneficial allele to reach quasi-fixation is similar to that for the dormant portion up until the average time in the seed bank approaches the number of active individuals in the population (i.e., 1,000), the threshold between weak and strong seed-bank effects (Fig. 4b). Surprisingly, the average length of time until a beneficial allele is quasi-fixed among actively reproducing individuals decreases once the average number of generations in the seed bank exceeds the strong seed-bank threshold, approaching the expected *T_fix_* for a population without a seed bank. This decrease in *T_fix_* is likely because the input of genetic diversity via resuscitation is low enough that it does not increase the amount of time required for a beneficial allele to reach fixation, a result that to our knowledge has not previously been reported. Contrary to the expectation that *T_fix_* increases with average time in the seed bank, our simulations suggest that under of a strong seed-bank effect, dormancy does not interfere with selection in the active portion of the population.

**Figure 4:**
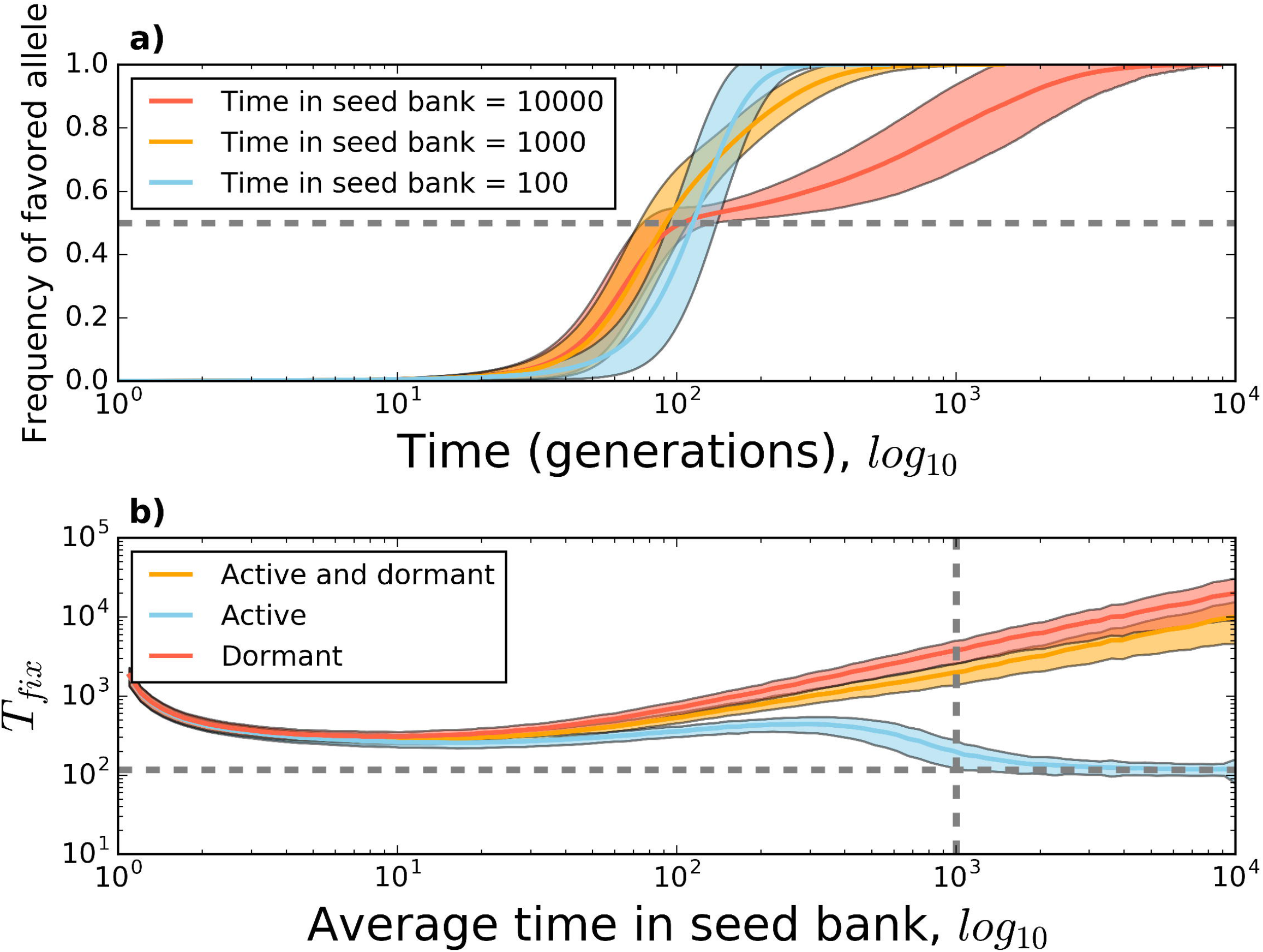
We performed population genetic simulations to determine the extent that dormancy alters the trajectory of a beneficial allele destined to go to fixation and the average length of time until it is fixed *T_fix_*. Simulations were performed on a population containing 1,000 active and 10,000 dormant individuals, where one active individual acquired a beneficial mutation with a 1% fitness advantage. **a)** The average length of time that an individual spends in a dormant state influences the trajectory of a beneficial allele destined to go to fixation. The horizontal grey line indicates an allele frequency of 0.5, showing how the beneficial allele first becomes fixed in the active portion of the population and requires an extended period of time to become fixed in the seed bank as the average length of time spent in a dormant state increases. b) The length of time until an allele is effectively fixed (*T_fix_*) for both active and dormant individuals scales with the average number of generations that an individual spends in a dormant state increases, we see that the trajectory of a beneficial allele destined for fixation changes shape. For example, after reaching an allele frequency of ~0.5 (horizontal grey dashed line), the rate that the beneficial allele increases in frequency drastically slows down when individuals remain dormant for 10,000 generations on average. b) The length of time until *T_fix_* for both active and dormant individuals scales with the average number of generations that an individual spends in a dormant state. However, *T_fix_* for the active portion of the population decreases once the average number of generations that an individual spends in a dormant state is higher than the number of active individuals in the population (vertical grey line). After this point, *T_fix_* for the active portion of the population quickly approaches *T_fix_* for an idealized population where the total population size is equal to the number of active individuals (horizontal grey line) (Eriksson et al., 2008, eq. 37). These results for **a)** and **b)** were obtained from a simulated Moran model of a selective sweep with a seed bank (see Supplementary Materials for details).

The decline of *T_fix_* for the active portion of a population subject to the strong seed-bank effect has important implications for the long-term evolutionary dynamics of microbial populations. Because the majority of a given bacterial genome is likely clonal, the rate of evolution in a population undergoing adaptive evolution can be sufficiently described by the beneficial mutation rate (*U*) and the strength of selection operating on a population of constant effective size (*N*) (Desai et al., 2007). Given that the average time it takes for a beneficial mutation to arise (*T_mut_*) is much greater than *T_fix_* the population will evolve at a rate (*v*), which is proportional to the rate that beneficial mutations of small effect arise and fix within a population (i.e., *v* ∝ *s^2^NU*) (Park et al., 2010). If this condition is violated, multiple beneficial mutations can arise on different genetic backgrounds and compete in the population, increasing *T_fix_* through the process of clonal interference (Gerrish & Lenski, 1998). As no one has examined clonal interference in an asexual population with a seed bank, there is no theory to draw on that would allow one to infer how dormancy would alter the rate of evolution in the multiple mutation regime. Our results suggest that the rate of evolution in the active portion of a microbial population with a strong seed-bank would not be pushed into the multiple mutation regime (Fig. 4b). However, *T_fix_* will increase if the average individual spends fewer generations in a dormant state than the number of active individuals (Fig. 4b). If the presence of a seed bank increases *T_fix_* to the point that it is not much larger than *T_mut_*, then multiple beneficial mutations will segregate simultaneously, moving the population into the multiple mutations regime. To fully describe the dynamics of clonal interference in a population with a seed bank it will be necessary to extend the single beneficial mutation dynamics described here to the multiple mutation regime using simulations and population genetic theory.

The ability to enter a dormant state has important implications for adaptive evolution. If seed banks increase the amount of genetic diversity, then it should also maintain beneficial alleles that allow for the population to rapidly adapt to new environments. It is often argued that microbial populations harbor low levels of genetic diversity across their genomes due to the effects of rapid linked positive selection (i.e., genetic draft) (Smith & Haigh, 1974; Gillepsie, 2000; Lynch, 2007). Under this scenario, the adaptive walk of a microbial population towards a fitness optimum consists of a series of origin-fixation steps (Orr, 2005), where one beneficial mutation goes to fixation before the next arises. Our simulations suggest that under a weak seed-bank effect the length of an adaptive walk (measured in generations) should increase proportionately with the average number of generations that an individual spends in a seed bank. However, under a strong seed-bank effect, the average *T_fix_* diverges for active and dormant individuals. This divergence suggests that the length of an adaptive walk will differ for active and dormant portions of the population, where the active portion of the population will reach the fitness peak in the same length of time as a population without a seed bank. All of these predictions assume that the adaptive walk consists of a series of sequential fixations of beneficial mutations. If multiple mutations segregate in the population at the same time, the length of the adaptive walk will increase. Whether dormancy increases or decreases the probability that the population is in the multiple mutation regime is a matter of ongoing research.

### Recombination

Dormancy can also preserve the diversity in microbial populations that is generated by recombination. The ability for microorganisms to acquire and exchange DNA by horizontal gene transfer (HGT) is a major evolutionary feature that can promote the spread of beneficial genes within a population (Medini et al., 2005; Lapierre & Gogarten, 2009; Dixit et al., 2016). The stochastic processes of gene gain and loss via HGT leads to the uneven distribution of genes among genomes in a population (Berg & Kurland, 2002; Baumdicker et al., 2012). This uneven distribution is often summarized as the total number of genes found within the population (i.e., the pangenome) and the number of genes shared among all individuals within the population (i.e., the core genome). Similar to how the presence of a seed bank increases the maximum amount of genetic diversity that can be maintained in a finite population (Fig. 3), it is possible that it would also increase the size of the pangenome. In addition, a strong seed-bank could alter the distribution of gene frequencies within a population, analogous to how the strong seed-bank effect alters the distribution of allele frequencies within a population (Blath et al., 2015). Thus, seed banks may alter the distribution of gene frequencies if inactive individuals are still capable of acquiring and incorporating foreign DNA into their genome. However, because the intake of DNA often requires the use of specific molecular systems (Chen & Dubnau, 2004; Mell & Redfield, 2014), the incorporation of foreign DNA into metabolically inactive cells is unlikely. Instead, it is more likely that the presence of a seed bank simply preserves genetic and genic diversity acquired by active individuals.

### Gene flow

Seed banks may affect microbial evolution by reducing the effect that migration has on allele frequencies within a population (i.e., gene flow). The ability to enter a dormant state can increase the chance that any two individuals share a common ancestor within a population (i.e., the probability of identity by descent) and reduce the amount of estimated differentiation among populations (i.e., the fixation index, *F_st_*) (Vitalis et al., 2004; Živković & Tellier, 2012; Hollander & Pederzani, 2017). While the effect of seed banks on the genetic similarity has been documented in plant populations (Tellier et al., 2011; Živković & Tellier, 2012), dormancy and migration are not always independent. For many organisms, dormancy can increase the probability of successfully migrating and establishing in a new population. For example, the spatial distribution of endospore-forming thermophilic bacteria in marine sediments closely reflects global ocean currents (Müller et al., 2014). However, dormancy and migration can represent a trade-off if the ability to enter a dormant state involves investing energy into a different mechanism than the one used to migrate (Olivieri, 2001). While this potential trade-off is well known in life history theory, it is necessary to examine it in a population genetic context in order to understand how dormancy and migration interact to shape patterns of shared genetic diversity between populations.

##### Box 1 Dormancy in low-energy environments

More than 50% of all prokaryotic (i.e., Bacteria and Archaea) cells in the oceans live underneath continents and on the sediment floor in a subsurface habitat known as the “deep biosphere” (Kallmeyer et al., 2012; Parkes et al., 2014). There, microorganisms are fueled by remnants of organic matter produced in the well-lit and productive surface waters that sink to the bottom of the ocean. As a result, the energy-limited microorganisms in the deep biosphere rest on the thermodynamic edge of life and death (Kallmeyer et al., 2012; Parkes et al., 2014; Jørgensen & Marshall, 2016; Starnawski et al., 2017). Because metabolic activity is extremely low and endospores are as abundant as vegetative cells, the deep biosphere likely constitutes the largest seed bank on the planet (Lomstein et al., 2012). Based on rates of amino acid racemization, it is estimated that the microbial biomass pool may only turnover once every thousand years, suggesting that populations within the deep biosphere are evolutionarily static (Lomstein et al., 2012; Hoehler & Jørgensen, 2013; Jørgensen & Marshall, 2016). Metagenomic and single-cell sequencing has revealed extremely low levels of genetic divergence for lineages within the deep biosphere (Starnawski, 2017). In addition, the ratio of synonymous to nonsynonymous polymorphisms did not change with sediment depth for three out of four lineages, suggesting that populations within the deep biosphere are not adapting to their environment (Starnawski et al., 2017). These findings are consistent with the expectation that dormancy acts as an evolutionary buffer. However, the slow turnover rate raises the question of whether dormancy is a viable life history strategy in the deep biosphere. Without a change in environmental conditions that would occasionally favor growth and reproduction, the deep biosphere may simply reflect a very large collection of microorganisms that are on a slow march to death.

In contrast, dormancy is likely a viable life history strategy in certain regions of the permafrost, a temperature and energy-limited environment covering 25% of Earth’s terrestrial surface (Graham et al., 2012). Upper layers of the permafrost that are not covered with ice can go through annual freeze-thaw cycles, providing the temporal variation necessary to favor dormancy as a life history strategy (Cohen, 1966; Malik & Smith, 2008). These freeze-thaw cycles could produce boom-and-bust periods of population growth that would leave a generation-time effect on the rate of molecular evolution, where lineages closer to the poles spend more time on average in a dormant state and show a reduced rate of molecular evolution. If so, this might result in latitudinal patterns of diversification, a long-standing pattern of diversity in a range of biological systems (Willig et al. 2003; Mittelbach et al. 2007). However, many microorganisms within the permafrost may not be completely dormant. For example, psychrophilic taxa isolated from the permafrost are capable of genome replication at temperatures as low as -20 °C (Tuorto et al., 2014) and evidence of DNA repair has been found in taxa that have persisted in the permafrost for hundreds of thousands to millions of years (Johnson et al., 2007; Dieser et al., 2013). This does not suggest that there are no dormant microorganisms within the permafrost, as the generation-time effect can still occur if growth rate is correlated with season and the energetic cost of DNA repair is thought to be negligible (see ***Mutation*** section). However, as average global temperature increases due to human-induced climate change, seasonal fluctuations below the freezing point of water will likely be less common. This rapid directional shift in the permafrost away from freeze-thaw cycles will likely impose a strong selective pressure against dormancy as a life history strategy, instead favoring sustained growth and reproduction.

## MACROEVOLUTIONARY CONSEQUENCES OF DORMANCY

In the previous section, we examined how seed banks influence the rate that allele frequencies change over relatively short periods of time within a lineage (i.e., microevolution). However, if seed banks buffer the aggregated effects of multiple evolutionary forces over long periods of time, then dormancy may have important implications for macroevolutionary phenomena. In the following section, we examine how dormancy influences macroevolutionary processes and patterns including the rate of molecular evolution, shared ancestry among lineages, speciation, and extinction.

### The rate of molecular evolution

If dormancy can affect the rate of evolution in microbial populations, we would likely observe it in bacteria that are capable to persist for extended periods of time in an inactive state via their capacity to form resistant, long-lived endospores. Many bacteria in the phylum *Firmicutes* possess the ability to form endospores, which are thought to be one of the most metabolically inert forms of life on the planet (Setlow, 2014). Low resource availability initiates endospore formation inside the mother cell. When development is complete, the mother cell is lysed and the endospore is released into the environment. (Tan & Ramamurthi, 2013). Endospores are persistent and in some instances, have reportedly been found to survive for hundreds of millions of years (Vreland, 2000). Endospores can endure exposure to extreme conditions such as high doses of gamma and UV radiation (Nicholson et al., 2000; 2005), desiccation (Nicholson et al., 2000), and the vacuum of space (Horneck et al., 1994; 2012). Given that endospore-forming bacteria can persist for long periods of time without reproducing, one might expect non-endospore-forming relatives to have more rapid rates of molecular evolution. This hypothesis was tested by analyzing a large collection of *Firmicutes* genomes, which included some isolates that had the ability to form endospores and other isolates that had lost the ability to form endospores (Weller & Wu, 2015). The rate of amino acid and synonymous substitutions for non-endospore forming isolates were significantly elevated relative to endospore-forming taxa and the phylogenetic branch length declined as the number of endospore-forming genes within a genome increased (Fig. 5a; Weller & Wu, 2015). These results suggest that rates of evolution increase when a lineage loses the ability to enter dormancy via sporulation. To further evaluate this effect of dormancy on the rate of evolution, we ran a simulation for the simple case of a neutrally evolving population, where we manipulated the average number of generations that an individual spends in the seed bank and calculated the substitution rate after 10,000 generations (see Supplementary Methods for more detail). For the entire population (i.e., both active and dormant individuals) we found that the rate of substitution declined as the average time in the seed bank increased, a finding that should apply to organisms across the tree of life irrespective of the mechanisms controlling transitions into and out of a dormant state (Fig. 5b).

**Figure 5:**
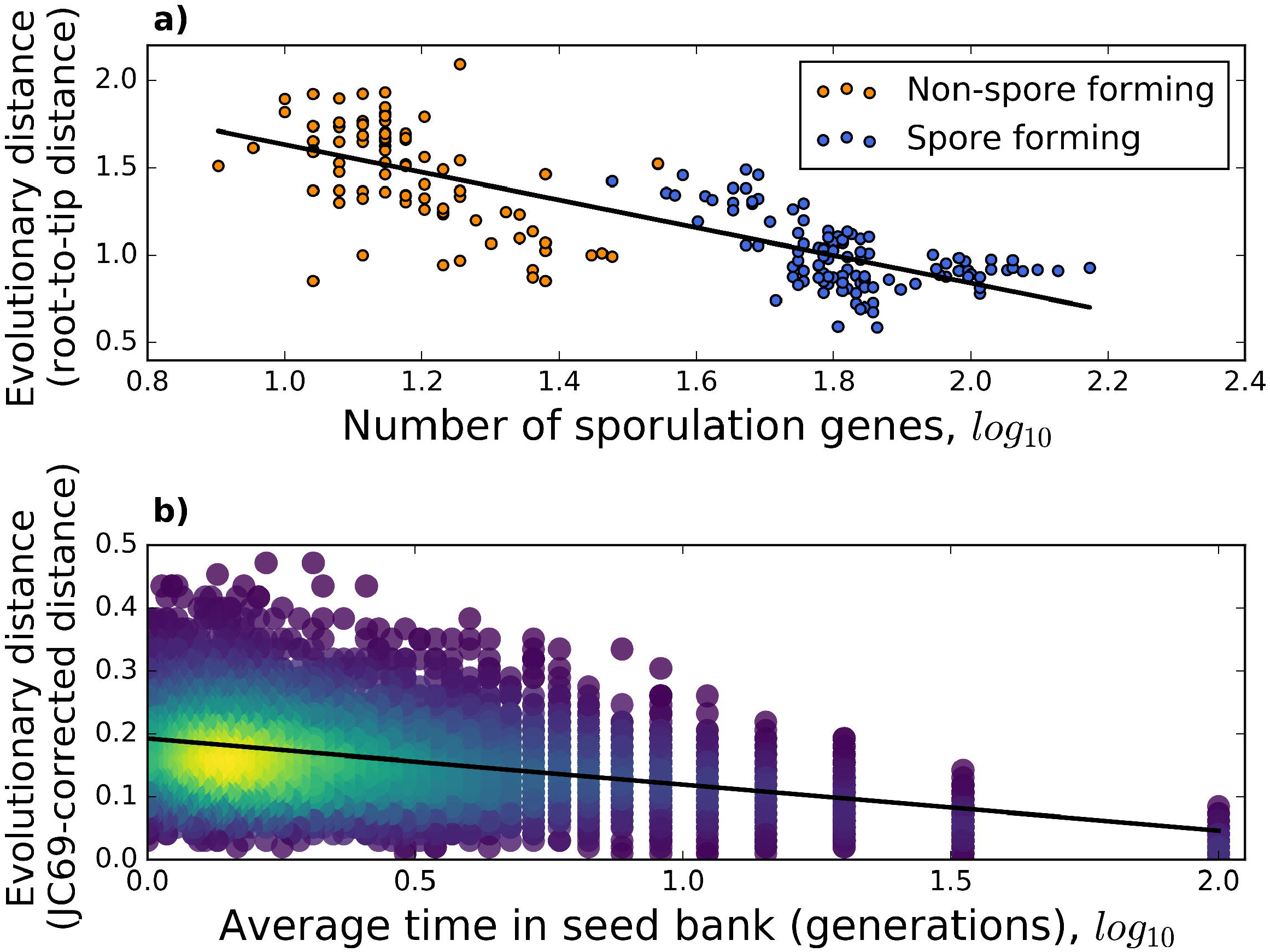
**a)** The evolutionary distance (calculated as the root-to-tip sum on the branch of the phylogeny) declines as the number of sporulation-associated genes increases among *Firmicutes* taxa, a common group of bacteria found in soils and hosts. The publically available data presented here is from the *Firmicutes* phylogeny of conserved genes (Weller & Wu, 2015). **b)** The evolutionary distance (calculated as the JC69 corrected distance) decreases as the average length of time that an individual spends in the seed bank increases (see Supplementary Materials). The lines in both **a)** and **b)** are the slopes from simple linear regression.

While theoretical and empirical evidence suggests that seed banks can reduce the rate of evolution within a lineage, this effect may be short lived if the ability to enter and exit a dormant state cannot be maintained. For example, the ability to form endospores has been lost multiple times within the *Firmicutes* phylogeny (Weller & Wu, 2015). The repeated loss of this seemingly beneficial trait across lineages can be explained by adaptive and non-adaptive evolutionary processes operating within a lineage. Organisms often need to invest in morphological or physiological structures to enter and exit a dormant state (Ayati & Klapper, 2012; Akiyama et al., 2017). The selective advantage dormancy must outweigh energetic costs as well as the fitness cost of not reproducing, where the strength and direction of natural selection can change over time (Simons, 2009). In relatively stable environments, the mechanism necessary to enter a dormant state may be lost from a lineage, either due to relaxed selection leading to the accumulation of mutations at dormancy-encoding loci or selection against the energetic cost of the mechanism (Dawes & Thornley, 1970). For example, in a long-term evolution experiment where five *Bacillus subtilis* populations were grown in culture medium that allowed for relaxed selection on endospore formation, all populations lost the ability to form endospores (Maughan et al., 2007). The accumulation of mutations due to relaxed selection was responsible for the loss of endospore formation in four of the five populations, while the loss of sporulation actually increased fitness in the remaining population. Endospore formation may be lost because it is a complex trait that is often encoded by more than 200 genes (Galperin et al., 2012), making it a large target for mutation. Alternatively, it is possible that the underlying genes may be lost because the energetic cost of even a single nucleotide is visible to natural selection in bacteria (Lynch & Marinov, 2015). In contrast to endospore-formation, other mechanisms regulating dormancy appear to require fewer genes than what is reported in endospore-forming bacteria. For example, resuscitation promoting factor (Rpf) is an extracellular enzyme encoded by a single gene that terminates dormancy for many bacteria belonging to the *Actinobacteria* phylum (Mukamolova et al., 1998; Keep et al., 2006). Rpf cleaves the 1,4-linkage between the amino sugars N-acetylglucosamine and N-acetylmuramic acid in peptidoglycan, which are major constituents in the cell walls of all bacteria. The release of Rpf by an actively growing individual into the environment wakes up neighboring cells, resuscitating them from a dormant state. The gene encoding Rpf likely has a lower probability of being lost owing to its small size and relatively low energetic cost. However, because it is released outside of the cell it can potentially wake up neighboring cells from different lineages, increasing the amount of competition for newly available resources. Ultimately, whether a mechanism used to enter a dormant state is retained in a lineage long enough to affect the rate of evolution will depend on the amount of environmental variation the lineage experiences over extended periods of time, its effect on fitness, the mutation rate of the lineage, the number of nucleotides that encode the mechanism, and the effective size of the population.

### Shared ancestry among lineages

Patterns of microbial genic ancestry can differ from patterns of species ancestry due to recombination (Degnan & Rosenberg, 2009). These dissimilar patterns of ancestry are often due to a few taxa harboring genes that have no detectable homologues in closely related lineages (i.e., ORFans) (Daubin & Ochman, 2004; Mira et al., 2004; Zhaxybayevaa et al., 2009). A popular explanation for the existence of ORFans is horizontal gene transfer (HGT). HGT is a reasonable hypothesis given that genic and genomic content can vary among and within microbial lineages due to the exchange of genetic material (Ochman et al., 2000). However, certain ORFans are essential for the process of cell division and it is unclear to what extent essential genes can be lost and acquired without resulting in the death of the cell.

To resolve this potential issue, the ability for microorganisms to remain in a dormant state for long periods of time has been proposed as an explanation for the existence of ORFans (González-Casanova et al., 2014). If some fraction of individuals can remain dormant for extremely long periods of time (González-Casanova et al., 2014), then the active portion of the population may continue to accumulate substitutions to the point that certain genes in the active pool no longer share homology with members in the dormant pool. To test this, a set of ORFan genes from the nitrogen-fixing soil bacterium *Azotobacter vinelandii* and neighboring lineages were examined (González-Casanova et al., 2014). These ORFan genes had the characteristic codon-usage and GC-content biases of *A. vinelandii*, but showed little homology to closely related members of the genus *Pseudomonas*, a result that was interpreted as support for the strong seed-bank hypothesis.

While the strong seed-bank hypothesis may help explain the long-term evolutionary consequences of remaining in a dormant state, an alternative hypothesis can explain the existence of ORFans. As a first step, additional copies of essential genes could be acquired by HGT, leading to relaxed selection on each gene copy. The original copy could then be physically lost due to gene deletion or functionally lost due to accumulated mutations (Lynch, 2007). This new copy would then gradually acquire the same GC-content as the rest of the genes within the lineage through a combination of lineage-specific mutation spectrum biases and the rate of GC-biased gene conversion (Lynch, 2007; Lassalle et al., 2015). In addition, lineage-specific selection on translational efficiency would shape the codon composition of the gene (Drummond & Wilke 2008; Plotkin & Kudla, 2011), particularly if the lineage has a large effective population size (Sharp & Li, 1987; Sharp et al., 2005). These directional pressures suggest that horizontally acquired genes that are retained in the genome may lose their signal of homology while becoming similar in composition to vertically acquired genes, a more parsimonious explanation than the strong seed-bank effect. While the theoretical models necessary to describe the effect of the seed bank on the genetic composition of microbial lineages exist (González-Casanova et al., 2014), further study is needed to determine whether the strong seed-bank effect applies to gene-specific patterns of ancestry.

### Speciation

While a microbial species definition has yet to be widely adopted (Rosselló-Móraa & Amann, 2015), population genetic processes can be used to infer how dormancy may alter the rate that microbial lineages diverge. Population genetic theory suggests that the presence of a seed bank can preserve genetic similarity among separate populations (Vitalis et al., 2004). The ability to exit a dormant state effectively acts like gene flow in physically separated populations, preserving genetic diversity and similarity that might otherwise be lost due to selection acting on newly arisen alleles within a population. The preserved genetic similarity due to the seed bank would likely slow the rate of evolutionary divergence between spatially separated lineages due to geographic divergence, reducing the rate of allopatric speciation. However, if a portion of the population remains in a dormant state while the rest of the population diverges to the extent that genetic material can no longer be exchanged, then dormant and active individuals may ultimately diverge into separate lineages. This process of divergence is an extension of the strong seed-bank effect previously discussed (González-Casanova et al., 2014), where the focus is on how evolutionary divergence between dormant and active individuals can prevent the exchange of genetic material, rather than on patterns of ancestry. Seed banks should influence the rate of speciation, but very little is known about how microbial lineages split or the rate that they split, regardless of the microbial species definition. While the rates of speciation are unclear, recent research on the net difference of speciation and extinction rates (i.e., diversification) in microorganisms suggests that microbial taxa split at a constant rate through time (Marin et al., 2017). If so, then we would expect that any decrease in the rate of speciation due to the presence of a seed bank would be associated with a similar increase in the rate of extinction, or vice-versa. The exact evolutionary mechanisms that are responsible for this constant rate of lineage diversification are currently unknown. Determining to what extent the ability to persist in a dormant state can alter the rate of speciation requires further research.

##### Box 2 Dormancy and the evolution of infectious diseases

The persistence and spread of pathogens requires that microorganisms contend with environmental variation inside and outside of their hosts. A host’s immune system represents a major challenge for pathogen survival. As a consequence, pathogens often exhibit periods of rapid growth during the early stages of infection, but slower growth in the later stages of infection and when individuals disperse outside of the host environment (Oliver et al., 1995). These temporal fluctuations in growth conditions suggest that the ability to enter a dormant state could be favored by selection. For example, *Yersinia pestis*, the causative agent of the plague, often enters a dormant stage that allows it to persist between infections (Pawlowski et al., 2011). In addition, *Y. pestis* contains several highly conserved genes that can contribute towards its ability to survive in a dormant state long after the host has expired (Easterday et al., 2012). Because the pathogen is capable of infecting a new host, any trade-off between virulence and transmission could potentially be reduced. If dormancy is as common among pathogens as it is in *Y. pestis*, then the ability to enter and exit a dormant state likely affects the evolutionary dynamics of pathogenic microorganisms.

Dormancy may also contribute to the evolution of antibiotic resistant pathogens. The evolution of disease-causing bacteria that are resistant to commonly used antibiotics is a major global health care concern (WHO, 2014). For example, methicillin-resistant *Staphylococcus aureus* infections alone are responsible for killing more than 11,000 US citizens each year (Gross, 2013; Golkar et al., 2014). By 2050, it is estimated that antibiotic resistant infections will have killed 10 million people (O’Neill, 2014). Because many antibiotics target molecular mechanisms that are primarily used during periods of growth, dormant cells are able of surviving antibiotic treatment, contributing towards the length and severity of antibiotic resistant infections (Wood et al., 2013; Zhang, 2014). Natural selection can optimize the length of time that bacteria remain dormant under antibiotic treatment (Fridman et al., 2014). In addition to preventing cell death, bacteria that survive antibiotic treatment by entering a dormant state are likely to evolve antibiotic resistance simply because they survive long enough to acquire a beneficial mutation (Levin-Reisman et al., 2017). To combat antibiotic resistant infections, it will be necessary to develop treatments and strategies to remove reservoirs of dormant bacteria. One potential strategy for targeting pathogens while minimizing the risk of antibiotic resistance is to resuscitate dormant cells alongside a course of antibiotics. The development of treatments designed to resuscitate dormant pathogenic bacteria is in the early stages, but holds promise. For example, commonly used clinical procedures were only able to detect *Mycobacterium tuberculosis* in sputum samples of infected individuals after cells were resuscitated with a dormancy-terminating extracellular enzyme, the resuscitating promoting factor (RPF; Mukamolova et al., 2010). However, because a large fraction of infections are acquired in hospitals (Klevens et al., 2007), it will be necessary to develop strategies to remove dormant disease-causing bacteria in buildings (i.e., the built environment) as well as in infected hosts. Recent work on characterizing the built environment suggests that the ability to persist shapes the composition of indoor microbial communities (Gibbons et al., 2015; Gibbons, 2016), a trait found in many disease-causing bacteria. Combating the emergence of antibiotic resistant bacteria and the role of dormancy as an adaptive trait will require re-examining the architecture and materials used to construct the buildings where infection is treated as well as infection treatment itself.

### Extinction

Because seed banks can reduce the average death rate in a population, they can likely buffer populations from extinction (Kalisz & McPeek, 1993). However, while next to nothing is known about extinction rates in microorganisms (Weinbauer & Rassoulzadegan, 2007), we can extend evolutionary models to examine how the ability to enter a dormant state might alter the rate of extinction. While ecological and environmental factors certainly contribute to the rate of extinction, we will focus on how evolutionary dynamics alter the probability of extinction for a microbial population. Because a large portion of bacterial genomes are thought to be clonal (Bobay et al., 2015) it has been argued that the rate of extinction in a microbial lineage is determined by the fixation rate of deleterious mutations; a process known as Muller’s Ratchet (Muller, 1964; Felsenstein, 1974). Once a deleterious mutation is fixed in a clonal population, it can only be removed by the fixation of a reverse mutation at the same site or compensated by the simultaneous acquisition and fixation of a high fitness mutation (i.e., stochastic tunneling; Iwasa et al., 2004). The negative fitness effect of deleterious substitutions accumulated over time can reduce reproductive output, leading to a positive feedback where deleterious mutations continue to accumulate and reproductive output further declines over time.

This feedback loop eventually results in the extinction of the lineage, a phenomenon known as a mutational meltdown (Gabriel et al., 1993; Lynch et al., 1993). Because lineages persisting in a dormant state likely fix mutations at a far lower rate and have a lower rate of evolution (Fig. 5), extinction via mutational meltdown is less likely to occur within lineages capable of forming a seed bank. Given that dormant individuals can persist on time scales upwards of millions of years (Cano et al., 1995; Greenblatt et al., 2004), it is worth investigating whether there is a relationship between the age of a microbial lineage and its ability to persist in a dormant state.

Analytical and conceptual models of microbial extinction have historically focused on clonal evolutionary dynamics. However, it is increasingly clear that microbes regularly exchange segments of DNA within and between lineages via HGT (Smith et al., 1993; Shapiro, 2016) and that the rate that DNA is exchanged between lineages can alter the rate of microbial diversification (Rayssiguier, et al., 1989; Dykhuizen & Green 1991; Fraser et al., 2007; Doroghazi & Buckley, 2014; Dixit et al., 2016; Marttinen & Hanage, 2017). The ability for a microorganisms to acquire and incorporate foreign DNA into their genomes suggests that fixed deleterious alleles can be purged from a population, alleviating the effects of Muller’s ratchet and mutational meltdown. This evolutionary scenario has been incorporated into analytical population genetic models, where populations capable of incorporating foreign DNA (whether from a donor cell or the environment) are effectively immune to Muller’s ratchet (Takeuchi et al., 2014). These results suggest that it is necessary for realistic models of microbial extinction to incorporate the per-base rate of recombination. However, even with an extremely high rate of recombination, we would still expect that the ability to enter a dormant state would reduce the probability of a lineage going extinct. Both HGT and dormancy can likely extend the lifespan of a microbial lineage in a fluctuating environment.

## CONCLUSION

The ability to enter a dormant state is a common life history strategy that is found across the tree of life. Historically dormancy has been viewed as an adaptive trait that preserves genetic and taxonomic diversity, with comparatively little attention given to its effects on the fundamental forces that underlie evolution. Using recent developments in theoretical population genetics and novel simulations we examined how the ability to enter a dormant state effects the population genetic forces that underlie evolutionary biology as well as common estimates of genetic diversity. Specifically, we identified the demographic scenarios where dormancy can affect natural selection. We then extended these results to determine how dormancy can influence the evolutionary dynamics of microbial lineages and in what environmental systems the ability to enter a dormant state is a viable life history strategy. We conclude that while dormancy can influence evolutionary dynamics and common estimates of genetic diversity, it is necessary to consider whether dormant organisms within a system are engaging in a life history strategy or simply reducing their level of metabolic activity out of physiological necessity. Other important, but yet unresolved questions include: 1) can dormancy push microbial populations into the multiple mutation regime, 2) how does dormancy as an environmentally dependent adaptive trait alter evolutionary dynamics and the process of adaptation, and 3) how has dormancy altered the rate of molecular evolution across the microbial tree of life. By considering the short-term population genetic and long-term macroevolutionary implications of dormancy as an adaptive trait, we can understand the extent that dormancy has impacted the evolutionary dynamics of the most abundant and metabolically diverse group of organisms on the planet.

Determining the extent that dormancy influences microbial evolution will require empirical research on whether dormant microorganisms in the environment are engaging in a life history strategy as well as continued development of the population genetic theory that accounts for the evolutionary effects of dormancy.

## ACKNOWLEDGMENTS

We acknowledge MG Behringer, J Davis, V Kuo, KJ Locey, RW Moger-Reischer, and NI Wisnoski for feedback on earlier versions of the manuscript. BK Lehmkuhl and E Polezhaeva provided technical assistance. This work was supported by National Science Foundation Dimensions of Biodiversity Grant 1442246 (JTL) and US Army Research Office Grant W911NF-14-1-0411 (JTL).

## DATA ARCHIVING STATEMENT

The data and code for the simulations used in this study can be found in a public GitHub repository (https://github.com/LennonLab/EvoDorm).

## AUTHOR CONTRIBUTIONS

WRS and JTL conceived of the ideas for the paper; WRS created and executed the simulations; WRS and JTL wrote the paper.

